# Do Goats Recognise Humans Cross-Modally?

**DOI:** 10.1101/2023.08.04.551944

**Authors:** Marianne A. Mason, Stuart Semple, Harry H. Marshall, Alan G. McElligott

## Abstract

Recognition plays a key role in the social lives of gregarious species, enabling animals to distinguish among social partners and tailor their behaviour accordingly. As domesticated animals regularly interact with humans, as well as members of their own species, we might expect mechanisms used to discriminate between conspecifics to also apply to humans. Given that goats can combine visual and vocal cues to recognize one another, we investigated whether this cross-modal recognition extends to discriminating among familiar humans. We presented 28 goats with facial photographs of familiar people and two repeated playbacks of a voice, either congruent (from the same person) or incongruent with that photograph (from a different person). When cues were incongruent, violating their expectations, we expected goats to respond faster and for longer after playbacks and show increases in physiological arousal (increased heart rate and/or decreased heart rate variability). We found the increase in latency that goats took to respond as the playback series progressed was greater when the face and voice were incongruent. As differences were not as predicted and only observed in one response measured, our evidence is tentative, but the variation in latency to look between congruency conditions suggests goat cross-modal recognition extends to humans. If this is the case, not only would this further demonstrate the flexibility of complex recognition systems to discriminate among members of a very different species, but indicates goats can produce mental representations for familiar people, a key component of individual recognition.

## 1. INTRODUCTION

Recognition forms a foundation for complex social behaviour, enabling animals to discriminate among social partners (e.g., between mates, kin and competitors) and tailor their behaviour accordingly (Tibbetts and Dale 2007; Wiley 2013; Yorzinski 2017). For domesticated animals, close-contact husbandry tasks (e.g., feeding, cleaning and health-checks) and repeated interactions (a prerequisite for recognition) with humans are a frequent occurrence. Through these interactions, dyadic social relationships can develop, requiring participants to recall outcomes of previous interactions with one other to anticipate the other party’s future behaviour (Hemsworth and Coleman 2011). This proximity to humans over thousands of generations has made the domestic environment a unique setting for the development of interspecific communication, potentially fostering more cognitively demanding forms of perception of human cues (Avarguès-Weber et al. 2013; MacHugh et al. 2017).

Dogs (*Canis lupus familiaris*), cats (*Felis catus*) and horses (*Equus caballus*) can combine cues in two sensory modalities (visual and vocal) not only to recognize members of their own species, but also familiar humans (Adachi et al. 2007; Proops et al. 2009; Taylor et al. 2011; Proops and McComb 2012; Takagi et al. 2019). Known as cross-modal recognition, such an ability infers the existence of a mental representation, for a conspecific or person, which can be used to compare against available cues (a prerequisite of individual recognition: Proops et al. 2009). This ability may enable these companion species to discriminate among people with greater accuracy and would be especially advantageous when cues in a particular modality are attenuated or unavailable (Ratcliffe et al. 2016). For example, visual features become less discernible at lower light intensities or over distance, so under these conditions, we may expect an animal to rely more on what they can feel, smell or hear than what they can see.

Unlike species domesticated for companionship or as working animals, livestock are more exclusively kept for their products, but likewise rely on us for food, shelter and other resources (MacHugh et al. 2017; Jardat and Lansade 2021). Although livestock may be expected to be under comparatively weaker selection to interpret human cues and communicate with us, they have already been shown to have an impressive repertoire of social skills to call upon when interacting with humans (for review, Jardat and Lansade 2021). Comparatively little is known about complex recognition of humans in livestock, although goats (*Capra hircus*) appear to use cross-modal recognition to distinguish among conspecific social partners (Pitcher et al. 2017).

Goats were among the first livestock species to be domesticated, approximately 10,500 years ago (MacHugh and Bradley 2001; MacHugh et al. 2017). At least compared to horses and other companion animals like dogs and cats, goats generally do not possess a close ‘working’ relationship with humans, being primarily domesticated for meat, milk and hair products (MacHugh and Bradley 2001; but see pack goats: Sutliff 2019). However, during its early domestication, this species appears to have undergone strong selection for tameness (Dou et al., 2023), a process that has been identified as being pivotal for the development of more advanced social cognition of human cues (Hare et al. 2005). Indeed, goats have been shown to read a variety of human cues, from attentional cues (Nawroth et al. 2015, 2016a, b; Nawroth and McElligott 2017) and facial expressions (frowning from smiling: Nawroth et al. 2018) to communicative gestures (pointing and tapping: Kaminski et al. 2005; Nawroth et al. 2015, 2020).

Goat cue use when recognising conspecific social partners spans multiple sensory modalities. When presented with a pen mate and a less familiar herd member and a call from one of the pair, goats were able to match playbacks to the original caller, turning towards them accordingly (Pitcher et al. 2017). In addition to cross-modal recognition, olfactory and vocal cues are important for early recognition between mothers and offspring (Poindron et al. 2007; Briefer and McElligott 2011; Perroux et al. 2022), with mothers responding to their own kids’ calls over those of other familiar kids up to 13 months post-weaning (Briefer et al. 2012). Coat colour also appears important for kids to recognize their mothers (Ruiz-Miranda 1993; Briefer and McElligott 2011) and goats are capable of discriminating group-members from non-group-members even when the head of the target animal is hidden (Keil et al. 2012). Despite the extensive research into how goats recognize conspecifics, no investigation to date has explored cues that they might use to recognize their next most important social partners, humans.

To investigate whether goats can recognise familiar people cross-modally, we presented subjects with a facial photograph and a voice which were either from the same person (so were congruent) or from different people (were incongruent). Following use of similar congruency paradigms in other species (e.g., Adachi et al. 2007; Takagi et al. 2019), we predicted that if goats can recognize humans cross-modally, their responsiveness would increase when visual and vocal cues were incongruent, reflecting a violation of their expectations. Specifically, following presentation of a voice incongruent with visual cues, goats were expected to respond faster and for longer, as well as exhibiting a faster heart rate and lower heart rate variability (associated with heightened physiological arousal: Briefer et al. 2015a; Baciadonna et al. 2020). In conducting this research, we aimed to determine whether goat ability to develop cognitive representations for known individuals, a building block of individual recognition, extends beyond their own species.

## 2. METHODS

### 2.1. Ethics Statement

Our stimuli collection from human participants and all experimental procedures were approved by the University of Roehampton Life Sciences Ethics Committee (Ref. LSC 19/ 280), with the latter being in line with ASAB guidelines for the use of animals in research (ASAB/ ABS 2020). All tests were non-invasive and lasted a maximum of seven minutes per subject for each trial.

### 2.2. Study Site & Sample Population

We conducted experiments between 8^th^ September - 19^th^ October 2020 and 19^th^ May - 16^th^ July 2021 at Buttercups Sanctuary for Goats (http://www.buttercups.org.uk/) in Kent, UK (51°13’15.7"N 0°33’05.1"E). During these periods, the sanctuary was open to visitors. Goats had daily access to a large outdoor area (approximately 3.5-4 acres) and were kept individually or in small groups within a large stable complex at night (mean pen size = 3.5m^2^). Throughout the day, animals had *ad libitum* access to hay, grass and water, and were supplemented with commercial concentrate according to age and condition. Our final sample comprised 28 adult goats (17 castrated males and 11 intact females) which were very habituated to human presence, were of various breeds (five Anglo Nubians, one British Alpine, one Golden Guernsey, one Old English Feral, 11 Pygmy goats, two Saanens, three Toggenburgs and four mixed breeds) and ages (mean age ± SD = 9.42 ± 3.58 years), and had resided at the sanctuary for over eight months (for detailed subject information, see Online Resource 1 Table 1).

**Table 1.** Predictors of time taken for goats to look at the photograph following playbacks of a familiar person’s voice (binomial GLMM). Results concerning the primary effect of interest, congruency x playback number interaction, shown in bold.

| Explanatory Variable | $\beta$ | S.E. | z-value | p-value |
| --- | --- | --- | --- | --- |
| Intercept | -5.555 | 1.559 |  |  |
| Congruency (I)<sup>a</sup> | -0.760 | 1.147 | -0.66 | 0.508 |
| Playback No. (2)<sup>b</sup> | 0.311 | 0.006 | 53.68 | <2x10<sup>-16</sup>**** |
| **Congruency (I)<sup>a</sup> x Playback No. (2)<sup>b</sup>** | **1.761** | **0.009** | **186.13** | **<2x10<sup>-16</sup>\*\*\*\*** |
| Year (2021)<sup>c</sup> | 2.543 | 1.335 | 1.91 | 0.057 |
| Goat Sex (M)<sup>d</sup> | -1.089 | 1.352 | -0.81 | 0.421 |
| Human Gender (M)<sup>d</sup> | 2.489 | 1.435 | 1.74 | 0.083 |
| Photograph ID (P)<sup>e</sup> | -0.831 | 1.211 | -0.69 | 0.493 |
| Voice ID (P)<sup>e</sup> | 0.554 | 1.183 | 0.47 | 0.640 |
| Preliminary Duration | -0.030 | 0.008 | -3.54 | 0.0004*** |

Key: I = Incongruent; M = Male; P = Preferred Person. Reference Categories: a = Congruent; b = Playback Number 1; c = 2020; d = Female; e = Non-preferred Person. \* $p < 0.05$ ; \*\* $p < 0.01$ ; \*\*\* $p < 0.001$ ; \*\*\*\* $p < 0.0001$

### 2.3. Stimuli Collection & Preparation

We selected goats which had been described by a particular caretaker, volunteer or frequent visitor (from whom stimuli were collected for the current experiment) as being particularly responsive to them. We assumed the reported preferential attention given to a particular person (hereafter known as a preferred person) would indicate goats were more familiar with their individual characteristics. We always tested goats with stimuli from one preferred person and one gender-matched, non-preferred caretaker to avoid potential reliance on gender as a cue and with the aim of ensuring subjects were familiar with all human stimuli presented.

In 2020, we collected photographs and voice samples from two male and three female caretakers. Photographs and voice samples for each person were collected in a single session at a two-meter distance, outdoors (as stimuli would normally be experienced by goats) and at the same location and time of day. This ensured better consistency in lighting and auditory environment for the photographs and recordings which subjects would experience in various combinations over their experimental trials. In 2021, we collected stimuli from five additional volunteers and frequent visitors to the sanctuary and one familiar staff member not involved with carrying out husbandry procedures (five females and one male) who had developed bonds with individual goats. For these, we took photographs and voice samples based on availability, meaning similarity in time of day could not be maintained, and location was sometimes shifted based on prevailing light conditions (otherwise conditions were as described above). Goats tested in 2020 were subject to photographic and vocal stimuli from pairs of caretakers, but in 2021, one of the two people goats experienced stimuli from (the preferred person) was not responsible for carrying out husbandry procedures (e.g., hoof trimming), which goats likely perceived as aversive. This could have affected the nuance of the relationship with their preferred person for goats tested in 2020 compared to those tested in 2021.

To collect visual stimuli, we asked each person to maintain a neutral expression and face the camera (Panasonic Lumix DMC-FZ45) before several frontal photographs of their head and shoulders were taken against a white background. Photographs were later edited so only the head and neck were visible, processed to improve clarity and brightness, before being blown up to A3 landscape size (slightly larger than natural head-size). We used images instead of their real-life counterparts to avoid unintentional cuing of subjects (Samhita & Gross, 2013) and to restrict cues available to only visual ones. Goats have previously been shown to be capable of discriminating details presented in both black and white (Nawroth et al. 2018) and, as used here colour photographs (Bellegarde et al. 2017).

For vocal recordings, we asked each person to say the phrase: “Hey, look over here,” several times in a manner they would normally use to address goats. We avoided using goats’ names and other potentially salient words, (e.g., food-related vocabulary) to test whether potential vocal recognition can be generalised based on vocal features, rather than being restricted to specific familiar words or phrases (Kriengwatana et al. 2015). Voice samples were recorded using a Sennheiser MKH 416 P48 directional microphone and a Marantz PMD-661 digital recorder (sampling rate: 48kHz, with an amplitude resolution of 16 bits in WAV format). We selected the clearest recording with the lowest background noise (mean recording length ± SD = 1.28s ± 0.35) and shifted the mean amplitude to 70dB to ensure consistency between playbacks and compiled these into a playback sequence using Praat v.6.1 (Boersma and Weenink 2019). Playback sequences comprised five seconds of silence before the first voice sample, followed by 10s of silence, a repeat of that sample, and 30s of silence. We measured goat responses in the 10s of silence immediately following each voice sample (hereafter known as the response period).

### 2.4. Experimental Enclosure

The test enclosure was constructed out of opaque metal agricultural fencing and barred metal hurdles in a large outdoor paddock that goats had ready access to throughout the day. In 2020, we constructed this away from areas experiencing greater visitor traffic to reduce disturbance. However, due to this location also being further away from areas more commonly frequented by goats, in 2021 we constructed the enclosure to similar dimensions, but moved it to ease subject transit to the experimental arena. The enclosure was divided into five sections (Fig. 1; Online Resource 1 Fig. 1). Goats entered through the preparation pen where we equipped them for experiments. Trials took place in the experimental arena. A corner of the arena (the holding pen) was sectioned off with hurdles and used both for training (see section 2.6.) and to prevent subjects from having visual access to experimenters during trials. We erected a semi-opaque green barrier around the fencing’s edge to prevent subjects from being able to see inside the preparation pen and outside the arena to reduce distractions and unintentional cuing from experimenters, visitors and other goats. Two Sony CX240E video cameras (frame rate = 25 FPS) were positioned at the front and back of the arena. The camera at the front was hidden under camouflage netting to prevent subjects attributing it as the sound source. We placed a target bucket filled with compressed hay next to the hurdle barrier, directly in front of the stimuli array. Goats were encouraged to approach and inspect this bucket as from this position they would have a clear view of the photograph, unimpeded by the barrier.

**Fig. 1.**
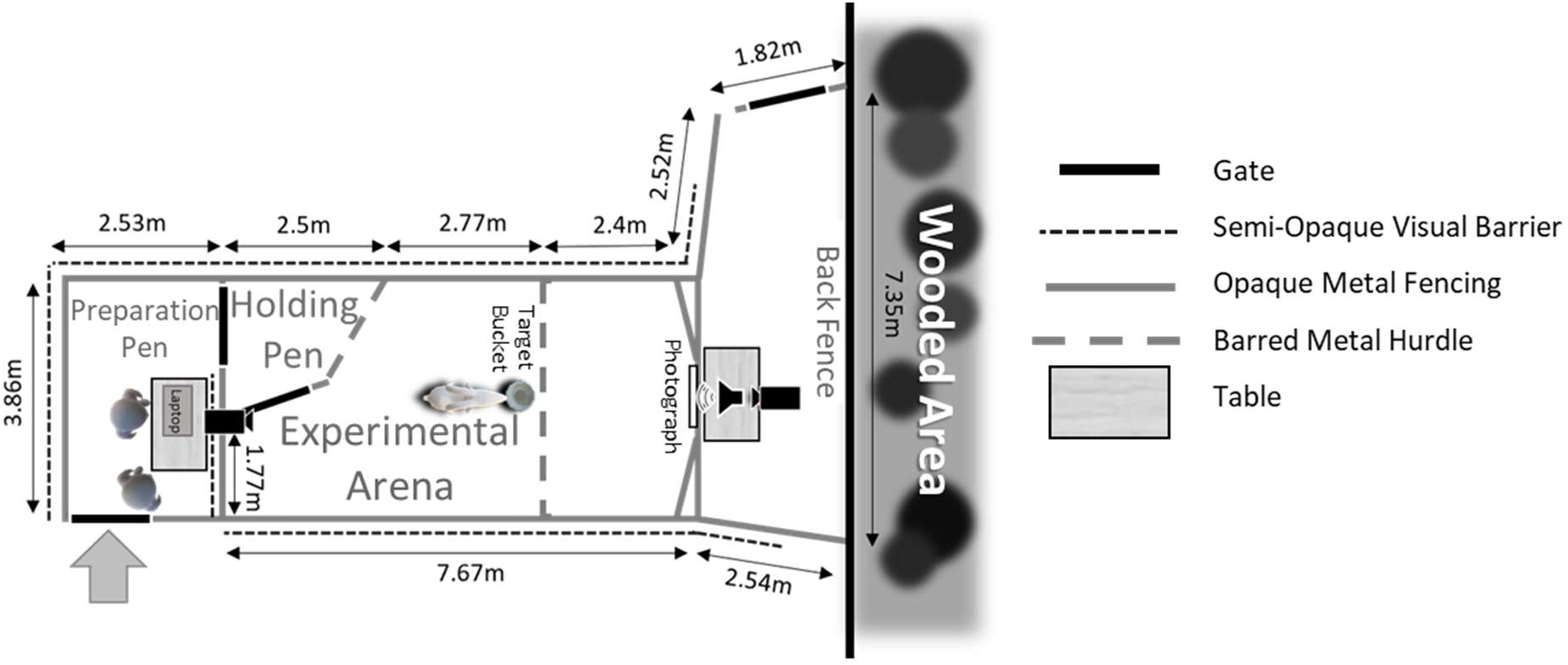
A schematic of the experimental enclosure used in 2020. Goats entered the enclosure through the preparation pen where they were fitted with a heart rate monitor, before being moved into the experimental arena for training and experimental trials.

During experiments, we affixed the appropriate photograph to a predetermined position at the front of the arena (top edge of image was 93.5cm off the ground). This position ensured that the speaker (a Bose Soundlink Mini Bluetooth Speaker II), which was placed directly behind the image was at approximately mouth level (70cm off the ground). The speaker used to broadcast playbacks in the current experiment has been verified to accurately reproduce the frequencies of the human voice (Ben-Aderet et al. 2017). The relative positioning of the speaker and photograph was aimed to direct subject attention towards the photograph and increase likelihood of goats perceptually coupling visual and vocal stimuli presented. We separated off the front of the arena using hurdles, imposing a distance between the subject and stimuli set of a minimum of 2.4m. This stopped goats from damaging photographs and was aimed to prevent them from localizing the speaker as the sound source. Goats have good visual acuity, being able to discriminate small details (3.4cm symbols) at distances of 1.5-2m (Blakeman and Friend 1986).

### 2.5. Habituation Phase

We habituated subjects to staying in the experimental arena alone for extended periods over three, four-minute sessions which took place prior to testing, and occurred between a minimum of three hours and a maximum of eight days apart (mean ± SD = 2 days 6 hrs ± 1 day 21 hrs). The variation in interval between successive habituation sessions was necessary as goats were not always willing to be led to the experimental enclosure or were displaced by more dominant individuals en route. Before starting each habituation session, we equipped goats with a Zephyr^TM^ BioHarness 3.0, which in conjunction with AcqKnowledge v.4.4.2 software (BIOPAC System Inc.) was used to transmit live cardiac data to a laptop (HP ProBook 650 G4) via Bluetooth during experimental trials. Similar wearable devices have been used to measure shifts in goat cardiac responses in relation to various factors (e.g., Briefer et al. 2015a, b; Baciadonna et al. 2016, 2019, 2020). If subjects exhibited signs of stress or aggression while this device was being fitted, we stopped and restricted future measurements taken during such trials to behaviour only (cardiac data collected from 16 out of 28 goats; for further details Online Resource 1 Table 1).

For the first habituation session, an experimenter sat still with their head down at the front of the arena where the photograph would be placed during experimental trials. As initially goats were not completely isolated, we aimed for this to reduce neophobia and encourage subjects to investigate their surroundings. For the following two habituation sessions subjects were kept alone in the arena. We occasionally repeated sessions when previous ones were halted due to adverse conditions such as rain, or when a long period of time had elapsed since the goat’s third habituation session (approximately seven days). For all three habituation sessions, we ensured animals had access to water and small food rewards (dry pasta), the latter of which were placed in and around the target bucket to encourage them to investigate and spend more time at the front of the arena. The habituation phase preceded the test phase by a minimum of three hours and 15 minutes and a maximum of eight days (median = 1 day; IQR = 1 day 4 hrs 38 mins).

### 2.6. Training Phase

The training phase took place directly before each experimental trial and aimed to incentivise goats to approach and investigate the target bucket during the test phase. Initially, goats entered the enclosure through the preparation pen where we fitted them with the BioHarness, which subjects had become accustomed to wearing during habituation sessions. To attain a clearer ECG trace, we clipped a patch of fur around the left shoulder blade at least one day prior to testing, over which the BioHarness module would be positioned. ECG gel was applied liberally to sensors on the BioHarness belt before it was placed around the subject’s thorax to improve conductance to the skin. Once a clear ECG trace had been secured, we led goats into the holding pen to begin training.

For training trial 1, Experimenter 1 and the subject were positioned inside the holding pen. Experimenter 2 held a piece of pasta (or cracker if goats were not sufficiently motivated by pasta) near the subject’s head and, once they had noticed it, walked slowly backwards towards the bucket while actively maintaining their attention. They then slowly and overtly placed two pieces of pasta inside, before crouching by the bucket and looking at it. Experimenter 1 then released the subject from the holding pen and stood aside, allowing the goat to pass and retrieve the pasta. The subject consumed both pieces of pasta before being led back into the holding pen. Training trial 2 proceeded similarly to the first, but instead of crouching down by the bucket, Experimenter 2 released the goat from the holding pen and stood aside to allow them to retrieve the reward. For training trials 3 and 4, Experimenter 2 gained the subject’s attention from the bucket, before baiting it and releasing them from the holding pen. Goats passed a trial if they successfully reached the bucket within 30s after release and needed to pass all four trials consecutively to proceed to the test phase. If a subject failed to pass a trial, the training process was repeated from the beginning and if they were not sufficiently motivated to participate in training, they were released, and the experimental trial was carried out on a different occasion.

### 2.7. Test Phase

Goats experienced four experimental trials each (3-14 days apart, mean = 7 days). Two trials were congruent (face and voice were from the same person) and two incongruent (from different people), with subjects experiencing all combinations of photograph and voice samples from a preferred person and a caretaker in a randomised order.

Once training had been completed, we led subjects back into the preparation pen and following checks to ensure the ECG signal had been maintained, they were distracted while the photograph was positioned. The goat was then led back into the arena and the holding pen shut behind them. Experimenters 1 and 2 quietly hid behind the visual barrier in the preparation pen and monitored subject movement in a nearby video camera’s LCD monitor (Fig. 1). We expected goats to approach and inspect the bucket (no pasta or water available during test phase) and when they were positioned close to and facing the front, we initiated playbacks (mean maximum amplitude ± SD = 76dB ± 2 measured 1m away under field conditions using a CEM^TM^ DT-8851 sound-level meter). If a subject moved away within the five seconds of silence preceding the first playback, where possible the playback sequence was aborted, and we re-initiated it when they were in a good position. If goats failed to be in a suitable position for six minutes following trial initiation (n = 11 trials), the trial was suspended, and they were released. We placed an event marker in the ECG trace to indicate occurrence of each playback for later analysis of cardiac measures. Goats that had been subject to both playbacks were released 30s after the final playback (median duration in arena = 1 min 20.95s; IQR = 64.12s). Of the 33 subjects tested, we excluded one from further testing due to health concerns, one for repeatedly failing to be in a suitable position for stimuli presentation, and a further subject and two trials from another due to a lack of training motivation.

### 2.8. Video Coding

We coded behavioural data using BORIS v.7.8.2 (Friard and Gamba 2016), with goat latency to look towards the photograph and looking duration in the 10s response periods following each playback defined to the nearest frame (0.04s). Behavioural measures were extracted from footage captured from the front and back of the arena, before being compared and combined. Measurements taken from the front were often clearer and without a blind spot, so these took precedence for quantifying behaviour. However, this was not always possible, with technical issues in the front camera for three trials and the back camera for two preventing experiments from being successfully captured by both, so in these cases we coded behaviours from the footage available. Under such circumstances, when goats went into the camera’s blind spot (n = 1 trial), only behaviours that could be coded with certainty were recorded, with others defined as missing.

Although we analysed behavioural data coded by a single observer, two further observers scored independent sets of videos comprising 37.9% and 42.1% of trials respectively (80% in total) to evaluate the reliability of our data set (Burghardt et al. 2012). Inter-observer reliability proved relatively high for duration (intraclass correlation analysis assuming Poisson distribution: R ± s.e.= 0.81 ± 0.03, *p*<0.0001, n = 1000 bootstraps; rptR R package: Stoffel et al. 2017) and latency measures (proportional distribution assumed: R ± s.e.= 0.93 ± 0.04, *p*<0.0001, n = 1000 bootstraps).

### 2.9. Exclusion Criteria

We applied two exclusion criteria when determining which trials should be considered for analysis. Firstly, if goats failed to look towards the stimuli array during both response periods in the playback sequence, the trial was excluded as subjects were interpreted as not being sufficiently attentive to human cues to notice incongruencies between them (n = 22 trials). Secondly, once the first exclusion criteria had been applied, we also excluded goats that did not look for at least one congruent and one incongruent trial as these prohibited within-subjects comparison (n = 2 subjects excluded). Ultimately, we analysed 95 trials from 28 subjects, nine of which experienced stimuli from men, and 19 from women.

### 2.10. Data Analysis

#### 2.10.1. General Model Parameters

We did not use automated model selection approaches for two reasons. Firstly, stepwise selection methods are known to lead to type 1 errors and the direction of selection (forwards or backwards) can have key implications for which variables get selected for the final model (Mundry and Nunn 2009). Second, even the currently favoured information criterion approaches (e.g., Akaike’s and Bayesian) may not perform substantially better (Bishop 2006; Mundry 2011; Forstmeier et al. 2017). Instead, as in addition to congruency, there were various factors which could have profound effects for goat behavioural and physiological responses we used a full model approach, as recommended by, for example, Forstmeier and Schielzeth (2011).

Our primary variable of interest for all models was congruency, i.e., whether the face and voice presented were from the same person (congruent), or from different people (incongruent). We also investigated whether goat responses changed over the playback series (playbacks 1 and 2), as could happen owing to, for example, habituation. In addition, we explored the interaction between congruency and playback number as there may be a lag before goats register and respond to incongruencies between stimuli presented. This interaction term was included in all models to begin with but removed if its effect was not significant to allow us to interpret the effects of playback number and congruency separately (as recommended by Engqvist 2005). In addition, as the experimental enclosure was erected at different locations in 2020 and 2021 and the pool of human stimuli collected was expanded in 2021, we controlled for the effect of year on goat responses. The goat’s sex and gender of the person whose identity cues were presented were also considered. This is because several studies have found sex-specific differences in responses to social stimuli (Proverbio 2017; Bognár et al. 2018), including in the context of cross-modal recognition (Proops and McComb 2012) and based on whether animals were experiencing stimuli from men or women (McComb et al. 2014; Shih et al. 2020; Li et al. 2020). Furthermore, we considered the identity of the photograph and voice, specifically whether or not they were derived from the preferred person, as this could have affected level of familiarity with or responsiveness to the cues presented. The interval between the experimenters leaving the arena (trial beginning) and the onset of playbacks (preliminary duration) was included as a covariate. Goats experiencing an extended time in the experimental enclosure may have had longer to habituate or become more aroused due to prolonged isolation from conspecifics, potentially affecting their behavioural and emotional responses (e.g., Siebert et al. 2011; Briefer et al. 2015a). Finally, we also fitted the covariate ‘measurement period’ to all models predicting cardiac responses. Noise in the ECG trace meant heart rate and heart rate variability (HRV) often could not be calculated over the entire response period. Measurement period therefore refers to the total time over which it was possible to calculate these responses. Given shorter measurement periods were likely the result of reduced ECG signal quality, this factor potentially represented an important control variable (e.g., Reefmann et al. 2009; Briefer et al. 2015b). Finally, trial number (1-4) was nested within subject identity and included as a random intercept to control for repeated measurements taken for each goat both within and between trials, as was the unique identifier given to the photograph and playback used for an experimental trial (the same stimuli were used over multiple trials).

These variables were used to model the two behavioural and two physiological responses of interest: looking duration and latency to look at the stimuli array following playbacks, as well as heart rate and HRV. All statistical analyses were conducted using R (version 4.2.3.: R Core Team 2023).

#### 2.10.2. Latency to Look Following Playbacks

Due to the bimodal nature of latency responses, with peaks at zero seconds (subjects were looking at onset of the response period) and 10s (subjects did not look throughout the response period), we analysed this variable using a binomial generalised linear mixed model (GLMM; glmer function, lme4 package: Bates et al. 2015). *Post hoc* tests were conducted using the emmeans package with Tukey’s corrections to account for multiple comparisons (Lenth 2021). The specific photograph and playback used were removed as random effects due to issues with model convergence which could have made the latency models less robust than those modelling other responses.

#### 2.10.3. Looking Duration Following Playbacks

As looking duration was restricted within the bounds of 0-10s, model residuals did not fit a normal error structure and a relatively small number of goats looked for entire 10s response period (median looking duration = 2.70s; IQR =4.27s), we considered approaches more typically applied to count data. Data was overdispersed and zero-inflated (verified using the DHARMa package: Hartig 2021), therefore using the package glmmTMB, we compared the fit of multiple models assuming different error distributions (Poisson, zero inflated-Poisson and negative binomial: Brooks et al. 2017). According to Akaike’s Information Criterion (AIC) we found that the best fit model had a zero-inflated negative binomial type 1 error structure (assumes variance increases linearly with the mean: Brooks et al. 2017). Such an approach generates both a conditional and a zero-inflated model, with the former predicting duration values greater than zero seconds using a log link, and the latter the probability of a zero observation (goats did not look) using a logit link. Preliminary duration and the random effect voice identity were removed due to issues with model convergence.

#### 2.10.4. Heart Rate & Heart Rate Variability

We calculated heart rate as beats per minute (BPM) and HRV as the root mean square of successive differences between heartbeats multiplied by 1000 (RMSSD). Changes in both these measures are thought to indicate shifts in physiological arousal and have been shown in relation to a variety of contexts in goats (e.g., Briefer et al., 2015a, b; Baciadonna et al., 2016, 2020). Baseline heart rate and HRV was calculated ideally in the 10s preceding the onset of the playback sequence. Trials began when both experimenters had left the arena and to reduce the effect of human manipulation on cardiac responses, baseline heart rate and HRV were only calculated following this, which for some observations (n = 13 trials) meant the measurement period was less than 10s. When noise present in the ECG trace restricted this period to less than five seconds, where possible (preliminary duration before onset of playbacks greater than 10s), we expanded the time frame for calculating this baseline up to 30s prior to the first playback until a measurement period of 10s could be achieved (n = 7 trials).

We examined the difference in heart rate and HRV calculated in the response periods following playbacks 1 and 2 compared to the baseline period (ΔHR = HR_pb1_ – HR_baseline_ or ΔHR = HR_pb2_ – HR_baseline_; ΔHRV = HRV_pb1_ – HRV_baseline_ or HRV_pb2_ – HRV_baseline_). These measures were used as variation in baseline heart rate and HRV meant relative changes in these responses measured over a single trial were more meaningful than absolute differences between individuals and trials (mean baseline BPM ± SD = 115.15 ± 16.97; mean baseline RMSSD ± SD = 22.42 ± 25.57).

Model residuals conformed to an approximately normal error structure, so we fitted a linear mixed model (LMM) to goat heart rate responses (lmer function, lme4 package: Bates et al. 2015). However, having fit a LMM to HRV data and visualised its residual variance, plots indicated the presence of extreme values. As these outliers had the potential to have a disproportionate effect when estimating model parameters, steps were taken to identify and exclude such observations. Using a z-score method (outliers package), we identified HRV values falling outside the 95% quantiles. After excluding these observations (n = 8), LMM assumptions were met so we used this approach to model shifts in HRV relative to congruency accordingly (lme4 package: Bates et al. 2015). To resolve convergence issues, we removed the specific photograph used as a random effect in models analysing HRV, and the photograph and playback used for those modelling heart rate responses.

## 3. RESULTS

### 3.1. Latency to Look Following Playbacks

Although goats in both the congruent and incongruent conditions took longer to respond to playback 2 than playback 1 (congruent playback 1 vs congruent playback 2: β ± s.e. = -0.311 ± 0.006, Z-ratio = -53.68, *p*<0.0001; incongruent playback 1 vs incongruent playback 2: β ± s.e. = -2.073 ± 0.007, Z-ratio = -277.31, *p*<0.0001), this increase in latency was greater in goats experiencing incongruent human cues (congruency x playback number interaction significant: Table 1; Fig. 2a; see Online Resource 1 Table 2 for further *Post hoc* comparisons). Additionally, goats spending a longer time in the arena before playbacks were initiated, tended to respond more quickly (Table 1). Goats also took a marginally longer time to respond to men’s versus women’s voices and in 2021 versus 2020. Goat sex, and whether the photograph and the voice presented were from a preferred person did not significantly affect their latency to look.

**Fig. 2.**
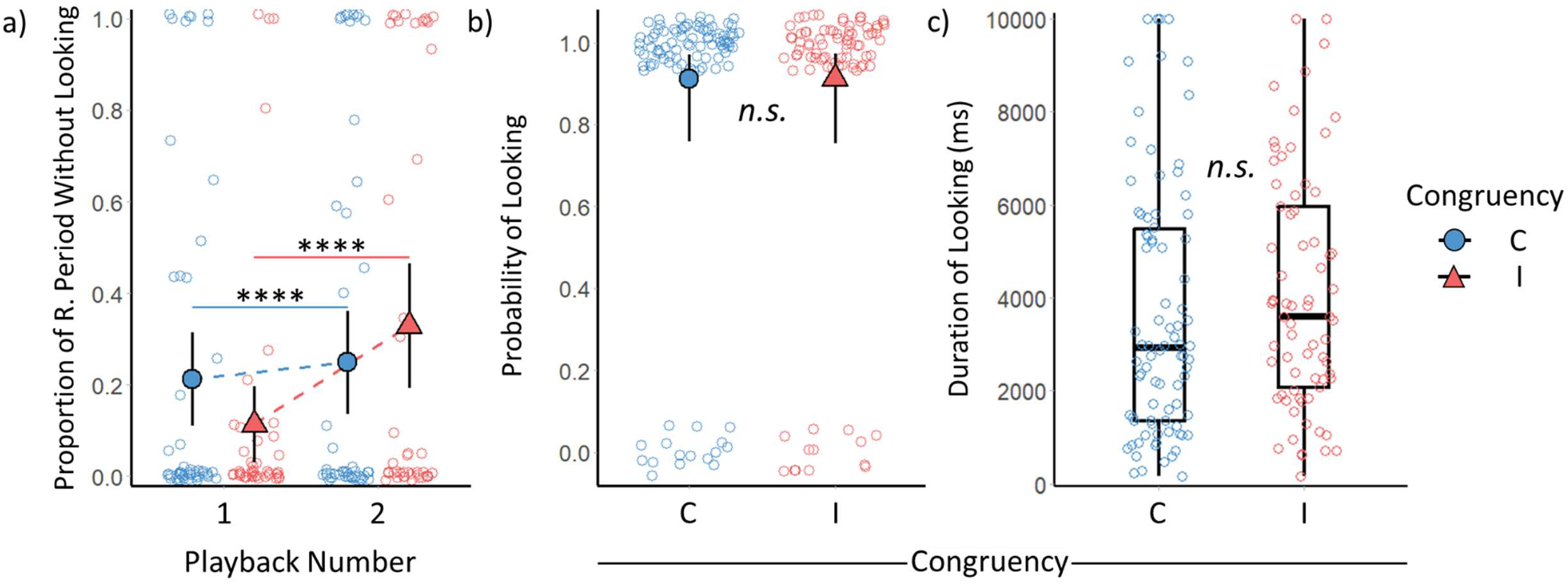
Effect of congruency between human facial and vocal cues on goat looking behaviours. a) The mean and confidence intervals for the effect of congruency and playback number on the proportion of the 10s response period that passed before goats looked. b) Predicted probability and 95% confidence intervals of goats looking in the congruent and incongruent conditions after playbacks. c) Median and interquartile range (IQR) for looking duration values greater than zero as a function of congruency. Boxplot whiskers extend to the maximum and minimum value less than 1.5 times above or below the IQR respectively. R. Period = Response period; C = Congruent; I = Incongruent. *n.s.* = non-significant; \*\*\*\**p*<0.0001

**Table 2.** Predictors of the likelihood that a goat did not look (zero-inflated model) and of their looking duration (conditional model) at the photograph following voice playbacks. Parameter estimates come from a zero-inflated negative binomial type 1 model. Results concerning the primary effect of interest, congruency, shown in bold.

| Explanatory Variable | Likelihood of Not Looking (zero-inflated model) |  |  |  | Looking Duration (conditional model) |  |  |  |
| --- | --- | --- | --- | --- | --- | --- | --- | --- |
| | $\beta$ | S.E. | z-value | p-value | $\beta$ | S.E. | z-value | p-value |
| Intercept | -2.219 | 0.606 |  |  | 8.170 | 0.170 |  |  |
| **Congruency (I)<sup>a</sup>** | **-0.030** | **0.419** | **-0.07** | **0.943** | **0.149** | **0.107** | **1.39** | **0.165** |
| Playback No. (2)<sup>b</sup> | 0.760 | 0.423 | 1.80 | 0.072 | -0.100 | 0.097 | -1.03 | 0.305 |
| Year (2021)<sup>c</sup> | 0.374 | 0.437 | 0.86 | 0.392 | -0.070 | 0.131 | -0.53 | 0.597 |
| Goat Sex (M)<sup>d</sup> | -0.176 | 0.435 | -0.41 | 0.685 | 0.165 | 0.133 | 1.24 | 0.216 |
| Human Gender (M)<sup>d</sup> | 0.163 | 0.444 | 0.37 | 0.714 | 0.091 | 0.137 | 0.66 | 0.509 |
| Photograph ID (P)<sup>e</sup> | 0.106 | 0.417 | 0.25 | 0.799 | -0.088 | 0.107 | -0.82 | 0.413 |
| Voice ID (P)<sup>e</sup> | -0.244 | 0.417 | -0.59 | 0.558 | -0.080 | 0.106 | -0.75 | 0.453 |

Key: I = Incongruent; M = Male; P = Preferred Person. Reference Categories: a = Congruent; b = Playback Number 1; c = 2020; d = Female; e = Non-Preferred Person.

### 3.2. Looking Duration Following Playbacks

How likely goats were to look following playbacks (Fig. 2b) or how long subjects looked when they did so (Fig. 2c) was not significantly affected by congruency between human facial and vocal identity cues (Table 2). However, goats were marginally more likely to look after playback 1 than playback 2. The year that goats were tested, their sex, the gender of the person providing stimuli and whether the face and voice belonged to the goat’s preferred person or not, also did not significantly affect how long or how likely they were to look after playbacks.

### 3.3. Heart Rate & Heart Rate Variability

Changes in goat heart rate (Fig. 3a) and HRV from baseline values (Fig. 3b) were not significantly affected by congruency between human identity cues (Table 3). However, goats did show a decrease in heart rate between playbacks 1 and 2 and had a marginally lower HRV when there was a longer measurement period available for calculating this response. There was no significant effect of playback number on HRV, or measurement period on heart rate, nor was there a significant effect of year the goat was tested, their sex, or the gender of the human stimuli experienced, whether these came from a preferred person, or the preliminary duration before onset of playbacks on heart rate or HRV.

**Fig. 3.**
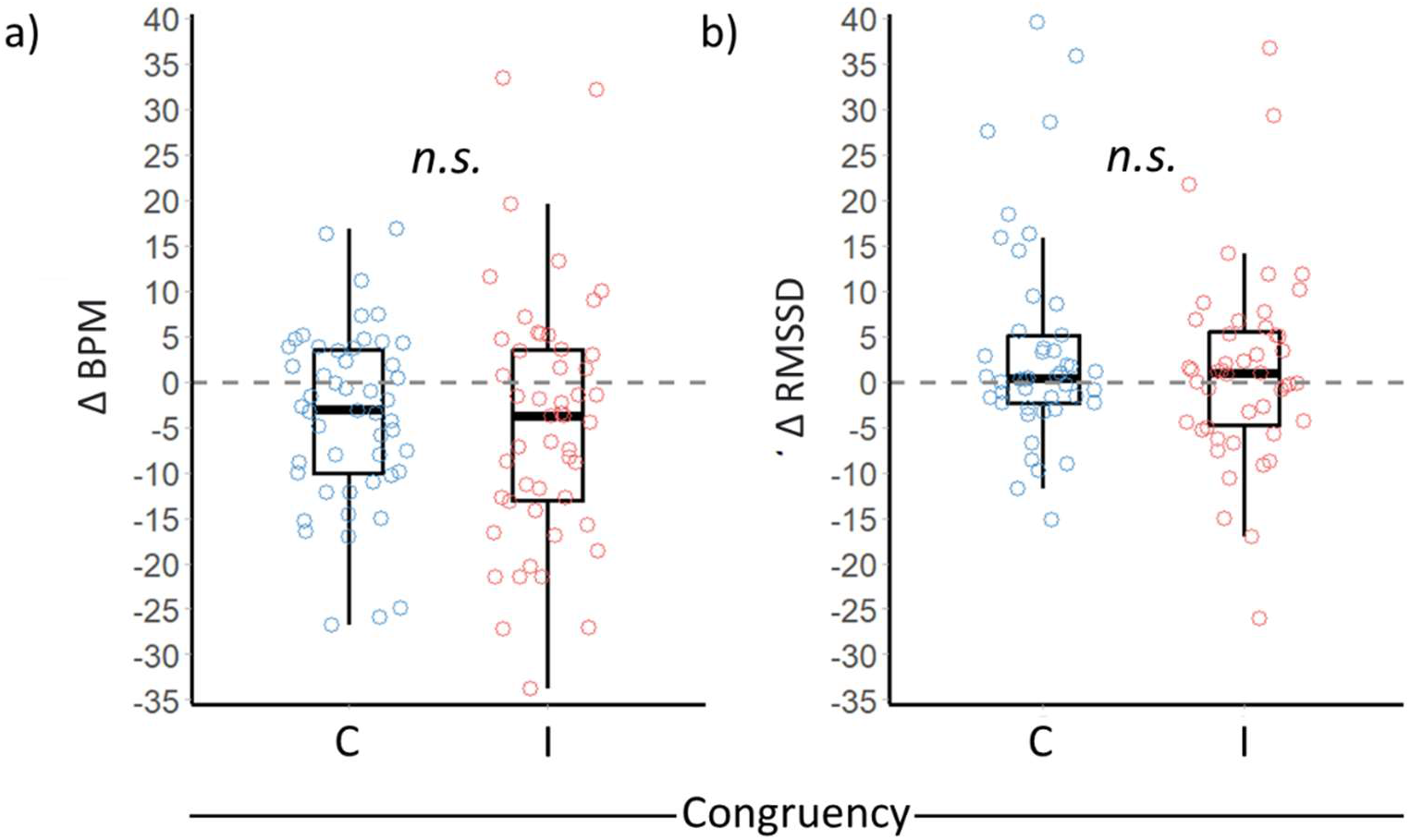
Effect of congruency between human visual and vocal identity cues on a) shifts in goat heart rate and b) HRV from baseline values (measured before onset of playbacks). Boxplots feature median and IQR. Whiskers extend to the maximum and minimum value less than 1.5 times above or below the IQR respectively. Baseline values are indicated by the dashed grey line. BPM = Beats per minute; C = Congruent; I = Incongruent. *n.s.* = non-significant

**Table 3.**
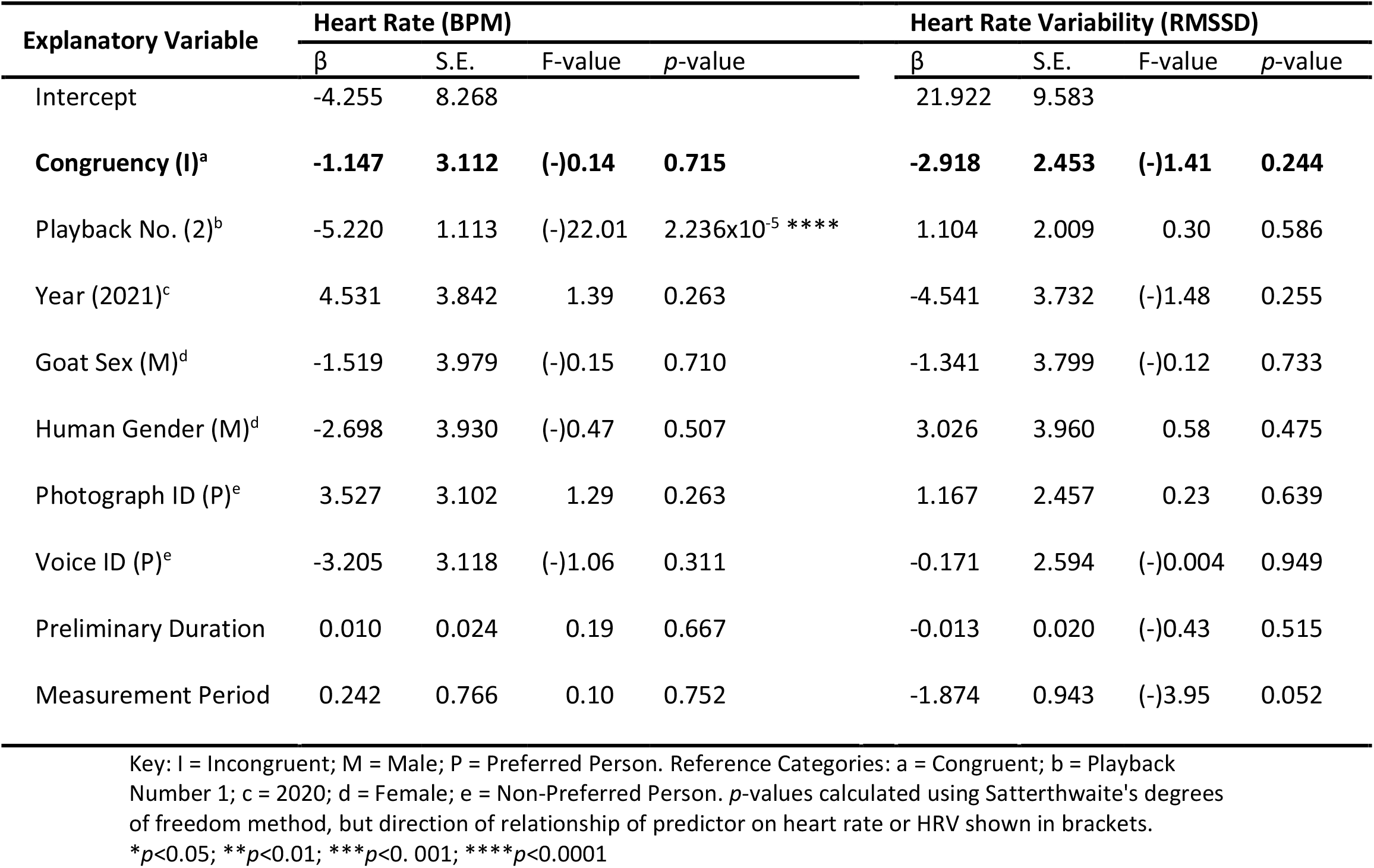
Predictors of goat heart rate and HRV, relative to baseline values (measured before onset of playbacks) in the response periods following presentation of a familiar human voice (LMM). Results concerning the primary effect of interest, congruency, shown in bold.

| Explanatory Variable | Heart Rate (BPM) |  |  |  | Heart Rate Variability (RMSSD) |  |  |  |
| --- | --- | --- | --- | --- | --- | --- | --- | --- |
| | $\beta$ | S.E. | F-value | *p*-value | $\beta$ | S.E. | F-value | *p*-value |
| Intercept | -4.255 | 8.268 |  |  | 21.922 | 9.583 |  |  |
| **Congruency (I)<sup>a</sup>** | **-1.147** | **3.112** | **(-)0.14** | **0.715** | **-2.918** | **2.453** | **(-)1.41** | **0.244** |
| Playback No. (2)<sup>b</sup> | -5.220 | 1.113 | (-)22.01 | 2.236x10<sup>-5</sup> **** | 1.104 | 2.009 | 0.30 | 0.586 |
| Year (2021)<sup>c</sup> | 4.531 | 3.842 | 1.39 | 0.263 | -4.541 | 3.732 | (-)1.48 | 0.255 |
| Goat Sex (M)<sup>d</sup> | -1.519 | 3.979 | (-)0.15 | 0.710 | -1.341 | 3.799 | (-)0.12 | 0.733 |
| Human Gender (M)<sup>d</sup> | -2.698 | 3.930 | (-)0.47 | 0.507 | 3.026 | 3.960 | 0.58 | 0.475 |
| Photograph ID (P)<sup>e</sup> | 3.527 | 3.102 | 1.29 | 0.263 | 1.167 | 2.457 | 0.23 | 0.639 |
| Voice ID (P)<sup>e</sup> | -3.205 | 3.118 | (-)1.06 | 0.311 | -0.171 | 2.594 | (-)0.004 | 0.949 |
| Preliminary Duration | 0.010 | 0.024 | 0.19 | 0.667 | -0.013 | 0.020 | (-)0.43 | 0.515 |
| Measurement Period | 0.242 | 0.766 | 0.10 | 0.752 | -1.874 | 0.943 | (-)3.95 | 0.052 |

Key: I = Incongruent; M = Male; P = Preferred Person. Reference Categories: a = Congruent; b = Playback Number 1; c = 2020; d = Female; e = Non-Preferred Person. *p*-values calculated using Satterthwaite's degrees of freedom method, but direction of relationship of predictor on heart rate or HRV shown in brackets. \**p*<0.05; \*\**p*<0.01; \*\*\**p*<0.001; \*\*\*\**p*<0.0001

## 4. DISCUSSION

We presented goats with a photograph of a familiar person and a voice that either matched (was congruent with) or did not match (was incongruent with) that photograph’s subject. If they could recognize the familiar person cross-modally, goats were predicted to respond faster and longer when there were incongruencies between stimuli, reflecting a violation of their expectations (e.g., Adachi et al. 2007; Takagi et al. 2019). Contrary to predictions, whether the photograph and voice were taken from the same or different people had no effect on how long or how likely goats were to look at the photograph following playbacks. Congruency between human facial and vocal cues also had no effect on goat heart rate or HRV, but did affect the latency it took for goats to look. Although this response also did not change in line with expectations, our findings suggest that goats had successfully perceived differences in congruency between the human visual and vocal information presented. In doing so, these results provide the first indication that like companion species and primates, goats may be able to use similarly complex cross-modal recognition systems to discriminate between human, as well as conspecific identity cues (Adachi et al. 2007; Sliwa et al. 2011; Proops and McComb 2012; Takagi et al. 2019).

Differences in time taken to look between trials with congruent and incongruent human identity cues suggest that goats were responding to changes in stimuli congruency. If so, they contribute to a growing body of literature emphasising the flexibility of complex recognition systems to create mental representations for members of a very different species (Ratcliffe et al. 2016). Within a species, recognition mechanisms are expected to co-evolve with signalling systems to facilitate communication between, for example, mates, kin, group members and competitors (e.g., Miller et al. 2020; Tibbetts et al. 2020). However, humans are a hugely important part of a goats’ life (and in those of other domesticated species) and as behaviour varies consistently between people, based on for example their attitudes, gender, skills and/ or experience (Hemsworth et al. 2000; Hemsworth and Coleman 2011; Ceballos et al. 2018; Celozzi et al. 2022), animals which discriminate between people may be favoured.

Robust recognition abilities like cross-modal recognition, which enable animals to discriminate among humans, as well as conspecifics, may allow better tracking of interactions with certain people, thereby forming a basis for interspecific social relationships (Tibbetts and Dale 2007; Hemsworth and Coleman 2011; Wiley 2013; Yorzinski 2017). Although isolated human-animal interactions often only have transient effects on animal emotional experiences and welfare, through the establishment human-animal relationships, repeated interactions can affect how animals perceive our cues in the long-term. Negative human-animal relationships, resulting from a negative perception of humans, have not only been linked to poor welfare, but can be detrimental to animal health and productivity, with fear of humans being the primary driving factor (reviewed by: Mota-Rojas et al. 2020). In contrast, in a positive human-animal relationship, social interactions with certain people may develop rewarding properties, providing opportunities for animals to experience positive emotions, a buffer against stressful situations (e.g., husbandry procedures) and potentially increasing an animal’s long term stress resilience (reviewed by: Rault et al. 2020). How well goats can discriminate among humans will affect whether experiences with certain people are attributed to that individual (individual recognition), people sharing similar features (class-level recognition, e.g., vets versus regular caretakers) or even just to humans in general (although recognition and generalisation are not mutually exclusive: Brajon et al. 2015; Yorzinski et al. 2017).

The evidence that goats registered incongruencies between human stimuli presented in different modalities could suggest the presence of an internal representation for familiar people; a foundation of individual recognition (Proops et al. 2009). However, as goats in the current study were required to discriminate cues from a single pair of familiar humans, we did not explicitly test whether cues they employed were at the individual (goats recognized both parties), or class-level (e.g., level of familiarity: Proops et al. 2009; Pitcher et al. 2017). Given the acknowledged importance of the human-animal relationship for welfare (Mota-Rojas et al. 2020; Rault et al. 2020), understanding recognition is important from such a perspective, as it affects the overall structure and complexity such relationships may take. Further research is needed to more fully understand the specificity and mechanisms that goats use to discriminate among people.

Goats in our experiment took longer to look as the playback series progressed, when human facial and vocal cues were incongruent. Conversely, similar congruency paradigms to the one used here have found a range of species tend to look quicker and/ or for longer when presented with incongruent conspecific cues (e.g., Proops et al. 2009; Gilfillan et al. 2016; Baciadonna et al. 2021) and, moreover, human identity cues in different modalities (e.g., Adachi et al. 2007; Takagi et al. 2019; Lampe and Andre 2012). Goats could have been quicker to respond to playback 1 (but not significantly so), but also habituated more quickly to these abnormal stimuli combinations, being slower to respond to playback 2 (again not significantly), together creating the observed difference in goat latency to look based on congruency. However, this cannot be currently confirmed and the reason why goats responded as they did remains unclear. Ultimately, although we observed changes in behaviour with stimuli congruency, these were not in line with expectations, nor was there any evidence of differences in physiological response between conditions.

We believe goat behaviour and physiology may not have changed as expected either due to factors related to goat social cognition or to our experimental design. Firstly, in order to register incongruencies, animals need to have developed an internal template for known individuals (Proops et al. 2009; Ratcliffe et al. 2016). Not all goats in our investigation may have possessed such a template for both people they were experiencing cues from, either through lack of cognitive ability or familiarity with their individual-specific cues. Secondly, we used photographs instead of live people. Photographs exclude olfactory, body (facial photographs were used), depth, perspective and motion cues and alter colour, all of which limits the amount of information goats would have had available to distinguish between individuals (Hill et al. 1997; Poindron et al. 2007; Fagot and Parron 2010; Keil et al. 2012; Lansade et al. 2020). Aside from it being more difficult for non-human animals to recognize objects from photographs, in order to have registered incongruencies between the visual and vocal information presented, goats would have also had to treat the photographs as representations of the people they depict (Fagot and Parron 2010). Indeed, a recent experiment found that goats did not express an immediate preference for photographs of group members over unknown conspecifics, nor did they learn at a faster rate when required to select a group member from three unknown individuals than vice versa (Langbein et al. 2023). These findings were interpreted as goats being unable to spontaneously link these photographs to their real-life counterpart, although it was suggested that they learnt to do this following presentation of different photographs of the same individual. Furthermore, the static, unresponsive nature of images can mean they are less salient and more rapidly habituated to than live stimuli (Vandenheede and Bouissou 1994; Bovet and Vauclair 2000).

In conclusion, our research provides evidence that goats may combine visual and vocal cues to recognize familiar humans, just as they can do with conspecifics (Pitcher et al. 2017). By extension, it suggests that goats may be able to form internal representations of heterospecifics, adding to a growing body of literature documenting the adaptability of complex cross-modal recognition systems to discriminate individuals of other, even phylogenetically very distant species (e.g., Adachi et al. 2007; Proops and McComb 2012). Overall, these findings may not only be important in furthering our basic knowledge of social cognition in human-animal relationships, but could also have vital applied implications for better understanding, and ultimately improving the welfare of domesticated animals.

## DATA AVAILABILITY

Data collected and R scripts used for the current study have been provided in the supplementary material (Online Resource 2).

## AUTHOR CONTRIBUTIONS

**Conceptualisation:** Marianne A. Mason, Stuart Semple, Alan G. McElligott; **Data curation:** Marianne A. Mason; **Formal analysis:** Marianne A. Mason, Harry H. Marshall; **Funding acquisition:** Marianne A. Mason, Stuart Semple, Alan G. McElligott; **Investigation:** Marianne A. Mason; **Methodology:** Marianne A. Mason, Stuart Semple, Alan G. McElligott; **Project administration:** Marianne A. Mason; **Supervision:** Stuart Semple, Harry H. Marshall, Alan G. McElligott; **Visualisation:** Marianne A. Mason, Harry H. Marshall; **Writing – Original draft:** Marianne A. Mason; **Writing – Review & editing:** Stuart Semple, Harry H. Marshall, Alan G. McElligott

## Supporting information

Subject information, photographs of the experimental enclosure and pairwise comparisons for the interaction between congruency and playback number

## ACKNOWLEDGEMENTS

We are very grateful to all of our field assistants and especially to Karen Lorena Estupinan Cely, Anastasia Makhlouf, Ellen Ye and Daniella Bernal Vega. We would also like to thank everyone who provided photographs and voice samples for our experiments, and to staff and volunteers at Buttercups Sanctuary for Goats for their help and advice and allowing us access to their goats.

## REFERENCES

Adachi I, Kuwahata H, Fujita K (2007) Dogs recall their owner’s face upon hearing the owner’s voice. Anim Cogn 10:17–21. https://doi.org/10.1007/s10071-006-0025-8

ASAB/ ABS (2020) Guidelines for the treatment of animals in behavioural research and teaching. Anim Behav 159:I–XI. https://doi.org/10.1016/j.anbehav.2019.11.002

Avarguès-Weber A, Dawson EH, Chittka L (2013) Mechanisms of social learning across species boundaries. J Zool 290:1–11. https://doi.org/10.1111/jzo.12015

Baciadonna L, Nawroth C, McElligott AG (2016) Judgement bias in goats (*Capra hircus*): Investigating the effects of human grooming. PeerJ 4:e2485. https://doi.org/10.7717/peerj.2485

Baciadonna L, Briefer EF, Favaro L, McElligott AG (2019) Goats distinguish between positive and negative emotion-linked vocalisations. Front Zool 16:25. https://doi.org/10.1186/s12983-019-0323-z

Baciadonna L, Briefer EF, McElligott AG (2020) Investigation of reward quality-related behaviour as a tool to assess emotions. Appl Anim Behav Sci 225:104968. https://doi.org/10.1016/j.applanim.2020.104968

Baciadonna L, Solvi C, La Cava S, Pilenga C, Gamba M, Favaro L (2021). Cross-modal individual recognition in the African penguin and the effect of partnership. Proc R Soc B 288:20211463. https://doi.org/10.1098/rspb.2021.1463

Bates D, Maechler M, Bolker B, Walker S. (2015) Fitting linear mixed-effects models using lme4. J Stat Softw 67:1–48. https://doi.org/10.18637/jss.v067.i01

Bellegarde LG, Haskell MJ, Duvaux-Ponter C, Weiss A, Boissy A, Erhard HW (2017). Face-based perception of emotions in dairy goats. Appl Anim Behav Sci 193:51–59. https://doi.org/10.1016/j.applanim.2017.03.014

Ben-Aderet T, Gallego-Abenza M, Reby D, Mathevon N (2017) Dog-directed speech: Why do we use it and do dogs pay attention to it? Proc R Soc B 284:20162429. https://doi.org/10.1098/rspb.2016.2429

Bishop CM (2006) Pattern recognition and machine learning. Springer Science + Business Media, New York.

Blakeman NE, Friend TH (1986) Visual discrimination at varying distances in Spanish goats. Appl Anim Behav Sci 16:279–283. https://doi.org/10.1016/0168-1591(86)90120-6

Boersma P, Weenink D (2019) Praat: Doing phonetics by computer. Version 6.1. retrieved 5^th^ June 2019 from http://www.praat.org/

Bognár Z, Iotchev IB, Kubinyi E (2018) Sex, skull length, breed, and age predict how dogs look at faces of humans and conspecifics. Anim Cogn 21:447–456. https://doi.org/10.1007/s10071-018-1180-4

Bovet D, Vauclair J (2000) Picture recognition in animals and humans. Behav Brain Res 109:143–165. https://doi.org/10.1016/S0166-4328(00)00146-7

Brajon S, Laforest JP, Bergeron R, Tallet C, Devillers N (2015) The perception of humans by piglets: Recognition of familiar handlers and generalisation to unfamiliar humans. Anim Cogn 18:1299–1316. https://doi.org/10.1007/s10071-015-0900-2

Briefer EF, McElligott AG (2011) Mutual mother-offspring vocal recognition in an ungulate hider species (*Capra hircus*). Anim Cogn 14:585–598. https://doi.org/10.1007/s10071-011-0396-3

Briefer EF, Padilla de la Torre M, McElligott AG (2012) Mother goats do not forget their kids’ calls. Proc R Soc B 279:3749–3755. https://doi.org/10.1098/rspb.2012.0986

Briefer EF, Tettamanti F, McElligott AG (2015a) Emotions in goats: Mapping physiological, behavioural and vocal profiles. Anim Behav 99:131–143. https://doi.org/10.1016/j.anbehav.2014.11.002

Briefer EF, Oxley JA, McElligott AG (2015b) Autonomic nervous system reactivity in a free-ranging mammal: Effects of dominance rank and personality. Anim Behav 110:121-132. https://doi.org/10.1016/j.anbehav.2015.09.022

Brooks ME, Kristensen K, van Benthem KJ, Magnusson A, Berg CW, Nielsen A, Skaug HJ, Machler M, Bolker BM (2017) glmmTMB balances speed and flexibility among packages for zero-inflated generalized linear mixed modelling. R Journal 9:378–400. https://doi.org/10.3929/ethz-b-000240890

Burghardt GM, Bartmess-LeVasseur JN, Browning SA, Morrison KE, Stec CL, Zachau CE, Freeberg TM (2012) Perspectives - Minimising observer bias in behavioural studies: A review and recommendations. Ethology 118:511–517. https://doi.org/10.1111/j.1439-0310.2012.02040.x

Ceballos MC, Sant’Anna AC, Boivin X, de Oliveira Costa F, Carvalhal MVDL, Paranhos da Costa MJR (2018) Impact of good practices of handling training on beef cattle welfare and stockpeople attitudes and behaviours. Livest Sci 216:24–31. https://doi.org/10.1016/j.livsci.2018.06.019

Celozzi S, Battini M, Prato-Previde E, Mattiello S (2022) Humans and goats: Improving knowledge for a better relationship. Animals 12:774. https://doi.org/10.3390/ani12060774

Dou M, Li M, Zheng Z, Chen Q, Wu Y, Wang J, Shan H, Wang F, Dai X, Li Y, Yang Z, Tan G, Luo F, Chen L, Shi YS, Wu JW, Luo X-J, Nanaei HA, Niyazbekova Z, Zhang G, Wang W, Zhao S, Zheng W, Wang X, Jiang Y (2023) A missense mutation in RRM1 contributes to animal tameness. Science Advances 9:eadf4068. https://doi.org/10.1126/sciadv.adf4068

Engqvist L (2005) The mistreatment of covariate interaction terms in linear model analyses of behavioural and evolutionary ecology studies. Anim Behav 70:967–971. https://doi.org/10.1016/j.anbehav.2005.01.016

Fagot J, Parron C (2010) Picture perception in birds: Perspective from primatologists. Comp Cogn Behav Rev 5:132–135. https://doi.org/10.3819/ccbr.2010.50007

Forstmeier W, Schielzeth H (2011) Cryptic multiple hypotheses testing in linear models: Overestimated effect sizes and the winner’s curse. Behav Ecol Sociobiol 65:47–55. https://doi.org/10.1007/s00265-010-1038-5

Forstmeier W, Wagenmakers EJ, Parker TH (2017) Detecting and avoiding likely false-positive findings - A practical guide. Bio Rev 92:1941–1968. https://doi.org/10.1111/brv.12315

Friard O, Gamba M (2016) BORIS: A free, versatile open-source event-logging software for video/ audio coding and live observations. Methods Ecol Evol 7:1325–1330. https://doi.org/10.1111/2041-210X.12584

Gilfillan G, Vitale J, McNutt JW, McComb K (2016) Cross-modal individual recognition in wild African lions. Biol Lett 12:20160323. https://doi.org/10.1098/rsbl.2016.0323

Hare B, Plyusnina I, Ignacio N, Schepina O, Stepika A, Wrangham R, Trut L (2005). Social cognitive evolution in captive foxes is a correlated by-product of experimental domestication. Curr Biol 15:226–230. https://doi.org/10.1016/j.cub.2005.01.040

Hartig (2021) DHARMa: Residual diagnostics for hierarchical (multi-level/ mixed) regression models. R package version 0.4.3. retrieved 27^th^ August 2021 from https://CRAN.R-project.org/package=DHARMa

Hemsworth PH, Coleman GJ, Barnett JL, Borg S (2000) Relationships between human-animal interactions and productivity of commercial dairy cows. J Anim Sci 78:2821–2831. https://doi.org/10.2527/2000.78112821x

Hemsworth PH, Coleman GJ (2011) Human-livestock interactions: The stockperson and the productivity of intensively farmed animals, 2nd edn. CAB International, Wallingford.

Hill H, Schyns PG, Akamatsu S (1997) Information and viewpoint dependence in face recognition. Cognition 62:201–222. https://doi.org/10.1016/S0010-0277(96)00785-8

Jardat P, Lansade L (2021) Cognition and the human-animal relationship: A review of the sociocognitive skills of domestic mammals toward humans. Anim Cogn 25:369–384. https://doi.org/10.1007/s10071-021-01557-6

Kaminski J, Riedel J, Call J, Tomasello M (2005) Domestic goats, *Capra hircus*, follow gaze direction and use social cues in an object choice task. Anim Behav 69:11–18. https://doi.org/10.1016/j.anbehav.2004.05.008

Keil NM, Imfeld-Mueller S, Aschwanden J, Wechsler B (2012) Are head cues necessary for goats (*Capra hircus*) in recognising group members? Anim Cogn 15:913–921. https://doi.org/10.1007/s10071-012-0518-6

Kriengwatana B, Escudero P, Ten Cate C (2015) Revisiting vocal perception in non-human animals: A review of vowel discrimination, speaker voice recognition, and speaker normalization. Front Psychol 5:1543. https://doi.org/10.3389/fpsyg.2014.01543

Lampe JF, Andre J (2012). Cross-modal recognition of human individuals in domestic horses (*Equus caballus*). Anim Cogn 15:623–630. https://doi.org/10.1007/s10071-012-0490-1

Langbein J, Moreno-Zambrano M, Siebert K (2023). How do goats “read” 2D-images of familiar and unfamiliar conspecifics? Front Psychol 14:1089566. https://doi.org/10.3389/fpsyg.2023.1089566

Lansade L, Colson V, Parias C, Trösch M, Reigner F, Calandreau L (2020) Female horses spontaneously identify a photograph of their keeper, last seen six months previously. Sci Rep 10:6302. https://doi.org/10.1038/s41598-020-62940-w

Lenth RV (2021) emmeans: Estimated marginal means, aka least-squares means. R package version 1.6.1.

Li Y, Wan Y, Zhang Y, Gong Z, Li Z (2020) Understanding how free-ranging cats interact with humans: A case study in China with management implications. Biol Conserv 249: 108690. https://doi.org/10.1016/j.biocon.2020.108690

MacHugh DE, Bradley DG (2001) Livestock genetic origins: Goats buck the trend. PNAS 98:5382–5384. https://doi.org/10.1073/pnas.111163198

MacHugh DE, Larson G, Orlando L (2017) Taming the past: Ancient DNA and the study of animal domestication. Annu Rev Anim Biosci 5:329–351. https://doi.org/10.1146/annurev-animal-022516-022747

McComb K, Shannon G, Sayialel KN, Moss C (2014) Elephants can determine ethnicity, gender, and age from acoustic cues in human voices. PNAS 111:5433–5438. https://doi.org/10.1073/pnas.1321543111

Miller SE, Sheehan MJ, Reeve HK (2020). Coevolution of cognitive abilities and identity signals in individual recognition systems. Phil Trans R Soc B 375:20190467. https://doi.org/10.1098/rstb.2019.0467

Mota-Rojas D, Broom DM, Orihuela A, Velarde A, Napolitano F, Alonso-Spilsbury M (2020) Effects of human-animal relationship on animal productivity and welfare. J Anim Behav Biometeorol 8:196–205. https://doi.org/10.31893/jabb.20026

Mundry R, Nunn CL (2009) Stepwise model fitting and statistical inference: Turning noise into signal pollution. Am Nat 173:119–123. https://doi.org/10.1086/593303

Mundry R (2011). Issues in information theory-based statistical inference - A commentary from a frequentist’s perspective. Behav Ecol Sociobiol 65:57–68. https://doi.org/10.1007/s00265-010-1040-y

Nawroth C, von Borell E, Langbein J (2015) ‘Goats that stare at men’: Dwarf goats alter their behaviour in response to human head orientation, but do not spontaneously use head direction as a cue in a food-related context. Anim Cogn 18:65–73. https://doi.org/10.1007/s10071-014-0777-5

Nawroth C, Brett JM, McElligott AG (2016a). Goats display audience-dependent human-directed gazing behaviour in a problem-solving task. Biol Lett 12:20160283. https://doi.org/10.1098/rsbl.2016.0283

Nawroth C, von Borell E, Langbein J (2016b) ‘Goats that stare at men’- revisited: Do dwarf dwarf goats alter their behaviour in response to eye visibility and head direction of a human? Anim Cogn 19:667–672. https://doi.org/10.1007/s10071-016-0957-6

Nawroth C, McElligott AG (2017) Human head orientation and eye visibility as indicators of attention for goats (*Capra hircus*). PeerJ 5:e3073. https://doi.org/10.7717/peerj.3073

Nawroth C, Albuquerque N, Savalli C, Single MS, McElligott AG (2018) Goats prefer positive human emotional facial expressions. R Soc Open Sci 5:180491. http://doi.org/10.1098/rsos.180491

Nawroth C, Martin ZM, McElligott AG (2020) Goats follow human pointing gestures in an object choice task. Front Psychol 11:915. https://doi.org/10.3389/fpsyg.2020.00915

Perroux TA, McElligott AG, Briefer EF (2022) Goat kid recognition of their mothers’ calls is not impacted by changes in fundamental frequency or formants. J Zool 318:297–307. https://doi.org/10.1111/jzo.13017

Pitcher BJ, Briefer EF, Baciadonna L, McElligott AG (2017) Cross-modal recognition of familiar conspecifics in goats. R Soc Open Sci 4:160346. https://doi.org/10.1098/rsos.160346

Poindron P, Terrazas A, Montes MDLLN, Serafín N, Hernández H (2007). Sensory and physiological determinants of maternal behaviour in the goat (*Capra hircus*). Horm Behav 52:99–105. https://doi.org/10.1016/j.yhbeh.2007.03.023

Proops L, McComb K, Reby D (2009) Cross-modal individual recognition in domestic horses (*Equus caballus*). PNAS 106:947–951. https://doi.org/10.1073/pnas.0809127105

Proops L, McComb K (2012) Cross-modal individual recognition in domestic horses (*Equus caballus)* extends to familiar humans. Proc R Soc B 279:3131–3138. https://doi.org/10.1098/rspb.2012.0626

Proverbio AM (2017) Sex differences in social cognition: The case of face processing. J Neurosci Res 95:222–234. https://doi.org/10.1002/jnr.23817

Ratcliffe VF, Taylor AM, Reby D (2016) Cross-modal correspondences in non-human mammal communication. Multisensory Research 29:49–91. https://doi.org/10.1163/22134808-00002509

Rault JL, Waiblinger S, Boivin X, Hemsworth P (2020) The power of a positive human-animal relationship for animal welfare. Front Vet Sci 7:590867. https://doi.org/10.3389/fvets.2020.590867

R Core Team (2023) R: A language and environment for statistical computing. R Foundation for Statistical Computing, Vienna, Austria. retrieved from https://www.R-project.org/

Reefmann N, Wechsler B, Gygax L (2009) Behavioural and physiological assessment of positive and negative emotion in sheep. Anim Behav 78:651–659. https://doi.org/10.1016/j.anbehav.2009.06.015

Ruiz-Miranda CR (1993) Use of pelage pigmentation in the recognition of mothers in a group by 2-to 4-month-old domestic goat kids. Appl Anim Behav Sci 36:317–326. https://doi.org/10.1016/0168-1591(93)90129-D

Samhita L, Gross HJ (2013) The “clever Hans phenomenon” revisited. Commun Integr Biol 6:e27122. https://doi.org/10.4161/cib.27122

Shih HY, Paterson M, Georgiou F, Phemshana NA, Phillips CJ (2020) Who is pulling the leash? Effects of human gender and dog sex on human-dog dyads when walking on-leash. Animals 10:1894. https://doi.org/10.3390/ani10101894

Siebert K, Langbein J, Schön PC, Tuchscherer A, Puppe B (2011) Degree of social isolation affects behavioural and vocal response patterns in dwarf goats (*Capra hircus*). Appl Anim Behav Sci 131:53–62. https://doi.org/10.1016/j.applanim.2011.01.003

Sliwa J, Duhamel JR, Pascalis O, Wirth S (2011). Spontaneous voice-face identity matching by rhesus monkeys for familiar conspecifics and humans. PNAS 108:1735–1740. https://doi.org/10.1073/pnas.1008169108

Stoffel MA, Nakagawa S, Schielzeth H (2017) rptR: Repeatability estimation and variance decomposition by generalized linear mixed-effects models. Methods Ecol Evol 8:1639–1644. https://doi.org/10.1111/2041-210X.12797

Sutliff DJ (2019) Pack goats in the Neolithic Middle East. Anthropozoologica 54:45–53. https://doi.org/10.5252/anthropozoologica2019v54a5

Takagi S, Arahori M, Chijiiwa H, Saito A, Kuroshima H, Fujita K (2019) Cats match voice and face: Cross-modal representation of humans in cats (*Felis catus*). Anim Cogn 22:901–906. https://doi.org/10.1007/s10071-019-01265-2

Taylor AM, Reby D, McComb K (2011) Cross modal perception of body size in domestic dogs (*Canis familiaris*). PLoS One 6:e17069. https://doi.org/10.1371/journal.pone.0017069

Tibbetts EA, Dale J (2007) Individual recognition: It is good to be different. Trends Ecol Evol 22:529–537. https://doi.org/10.1016/j.tree.2007.09.001

Tibbetts EA, Liu M, Laub EC, Shen SF (2020) Complex signals alter recognition accuracy and conspecific acceptance thresholds. Phil Trans R Soc B 375:20190482. https://doi.org/10.1098/rstb.2019.0482

Vandenheede M, Bouissou MF (1994) Fear reactions of ewes to photographic images. Behav Proc 32:17–28. https://doi.org/10.1016/0376-6357(94)90024-8

Wiley RH (2013) Specificity and multiplicity in the recognition of individuals: Implications for the evolution of social behaviour. Biol Rev 88:179–195. https://doi.org/10.1111/j.1469-185X.2012.00246.x

Yorzinski JL (2017) The cognitive basis of individual recognition. Curr Opin Behav Sci 16:53–57. https://doi.org/10.1016/j.cobeha.2017.03.009

