## Supplementary material for "Do Goats Recognise Humans Cross-Modally?": Subject information, photographs of the experimental enclosure and pairwise comparisons for the interaction between congruency and playback number

### Do Goats Recognise Humans Cross-Modally? Supplementary Information

**Table 1.** Subject name, year in which they were tested, sex (M = Male; F = Female), breed, age, number of years at study site, order of trials included in analysis (C = Congruent, I = Incongruent) and whether cardiac, as well as behavioural data had been successfully taken during experimental trials (Y = Yes; N = No).

| Goat ID | Tested | Sex | Breed | Age (years) | Duration at Sanctuary (years) | Trial Sequence | Contributed Cardiac Data? |
|---|---|---|---|---|---|---|---|
| Bernard | 2020 | M | Anglo Nubian | 9 | 9 | C, C, I, I | Y |
| Davey | 2020 | M | Saanen | 15 | 7 | C, I, C, I | Y |
| Dixie | 2020 | F | Pygmy | 11 | 9 | C, I, C | N |
| Dylan | 2020 | M | Pygmy | 7 | 5 | C, I, I, C | N |
| Ewok | 2020 | M | Anglo Nubian | 4 | 3 | I, I, C, C | Y |
| Juliet | 2020 | F | Toggenburg | 8 | 3 | C, C, I, I | N |
| Luke | 2020 | M | Anglo Nubian | 4 | 3 | C, I, C, I | Y |
| Natalie | 2020 | F | Swiss-Alpine Cross | 10 | 9 | I, C, I | Y |
| Nigel | 2020 | M | Pygmy | 9 | 0.75 | I, I, C | N |
| Princess | 2020 | F | Anglo Nubian | 4 | 3 | I, C, C, I | Y |
| Spice | 2020 | F | Anglo Nubian | 7-8 | 3 | C, I | N |
| Tarnie | 2020 | F | Golden Guernsey | 16 | 4 | I, C, I, C | Y |
| Bill | 2021 | M | Toggenburg | ≈16 | 4 | I, C, I | Y |
| Bramble | 2021 | M | Pygmy | Unknown | ≈3 | I, I, C, C | Y |
| Cole | 2021 | M | Mixed Breed | ≈8 | 2 | C, I, C | N |

| | | | | | | | |
|---|---|---|---|---|---|---|---|
| Dunstan | 2021 | M | Pygmy | 9 | 7 | I, C, C, I | N |
| Franky | 2021 | M | Saanen Cross | 5 | 3 | I, C, I, C | Y |
| Glenda | 2021 | F | Saanen Cross | ≈10 | ≈1 | I, C, C | N |
| Goesover | 2021 | M | Saanen | Unknown | 1 | C, I, C | Y |
| Heidi | 2021 | F | Toggenburg | 11 | 9 | I, C, C | Y |
| Jet | 2021 | M | Pygmy | 9 | 4 | I, C | Y |
| Joseph | 2021 | M | Pygmy | 7+ | 6 | C, I, C | N |
| Khan | 2021 | M | Pygmy | 9+ | 4 | C, C, I, I | N |
| Kirk | 2021 | M | Pygmy | 9+ | 4 | C, I, C, I | Y |
| Mary | 2021 | F | British Alpine | 16+ | 4 | C, I, C | Y |
| Milly | 2021 | F | Old English Feral | Unknown | ≈4 | C, I, C | N |
| Pooky | 2021 | F | Pygmy | 9 | 5 | C, C, I | N |
| Sundance | 2021 | M | Pygmy | 13 | 7 | I, C, C | Y |

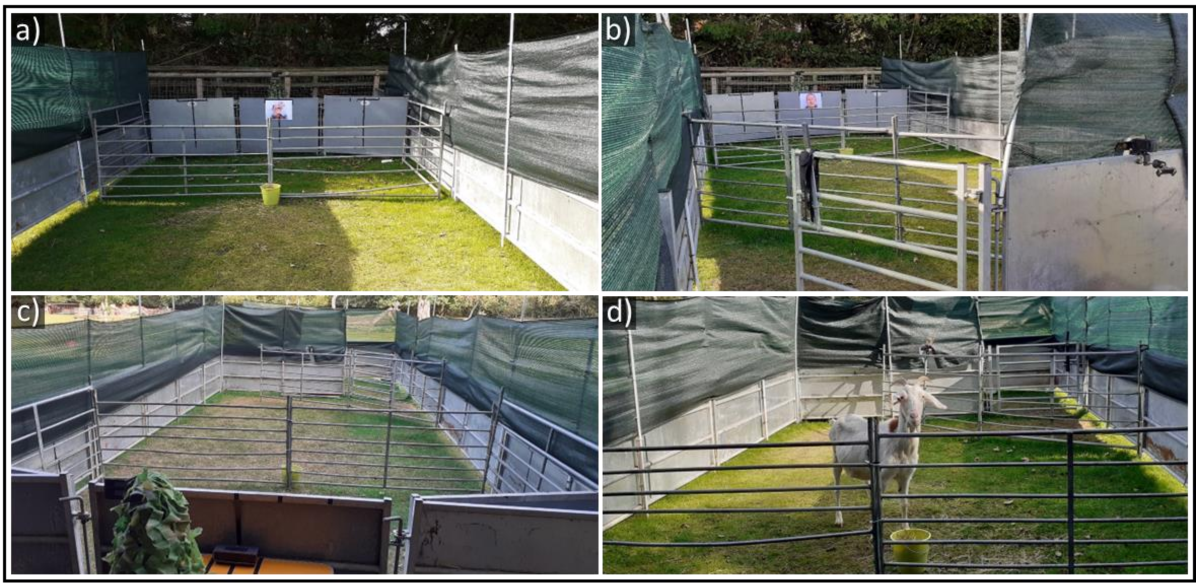


**Fig. 1.** a) A photograph showing the view of the front of the arena in 2020. The target bucket used to manoeuvre goats into a suitable position for stimuli presentation was placed directly in front of the photograph and speaker behind the front separator. b) A view of the front of the arena in 2020, including the holding pen and video camera positions. c) A view of the back of the arena, including the video camera (hidden under camouflage netting) and speaker position. d) This photograph shows a subject in an ideal position for stimuli presentation.

**Table 2.** Goat latency to look at the photograph following voice playbacks in relation to the effect of congruency between human visual and vocal cues, in combination with playback number (results of *post hoc* tests for congruency x playback number interaction). Significant pairwise comparisons are shown in bold.

| Explanatory Variable | B | S.E. | z-ratio | *p*-value |
|---|---|---|---|---|
| • **C1 - C2** | **-0.311** | **0.006** | **-53.68** | **<0.0001****** |
| • **I1 - I2** | **-2.073** | **0.007** | **-277.31** | **<0.0001****** |
| • C1 - I1 | 0.760 | 1.147 | 0.66 | 0.911 |
| • C1 - I2 | -1.313 | 1.147 | -1.14 | 0.662 |
| • I1 – C2 | -1.072 | 1.147 | -0.93 | 0.787 |
| • C2 – I2 | -1.001 | 1.147 | -0.87 | 0.819 |

Key: C1= Congruent condition, Playback 1; C2= Congruent Condition, Playback 2; I1= Incongruent Condition, Playback 1; I2= Incongruent Condition, Playback 2. ****$p<0.0001$
